# Shear Stress Initiates Endothelial to Mesenchymal Transition in Endocardial Endothelial Cells

**DOI:** 10.1101/2022.12.05.519049

**Authors:** Kathleen N. Brown, Kim T. Phan Hong, Tasneem Mustafa, Elysa Jui, Fariha N. Ahmad, Ravi K. Birla, Philippe Sucosky, Jennifer P. Connell, Sundeep G. Keswani, K. Jane Grande-Allen

## Abstract

Discrete subaortic stenosis (DSS) is a congenital heart disease in which a fibrotic membrane forms below the aortic valve; the underlying cellular mechanisms are currently unknown. Since an elevated pressure gradient in the left ventricular outflow tract (LVOT) is a distinguishing feature of DSS, it is hypothesized that the membrane formation is caused by elevated wall shear stress applied to the endocardial endothelial cells (EECs) that line the LVOT, triggering fibrosis. To correlate shear stress to an EEC fibrotic phenotype, we applied fluid shear stress to EECs at physiological and pathological shear rates using a cone-and-plate device, designed to recapitulate physiological wall shear stress in a controlled in vitro environment. Controlled shear stress regimes were applied to EECs to replicate the conditions observed in DSS patients. We found that elevated shear stress triggered EEC alignment as well as endothelial-to-mesenchymal transformation (EndMT) signaling pathways driven by upregulation of *SNAI1* gene expression. The EECs were then treated with a small molecule inhibitor of Snail1 protein, CYD19, to attempt to attenuate EndMT signaling, and subsequently subjected to pathological shear stress. The Snail1 inhibitor did downregulate selected markers of EndMT signaling, although only transiently. Interestingly, the application of shear stress had a greater effect on the EEC gene and protein expression than did the Snail1 inhibition. This investigation of EEC response to shear stress reveals the pronounced and complex effect of this mechanical stimulation on the EEC phenotype. Further study should reveal the mechanisms that drive fibrosis and the formation of the DSS membrane.

## Introduction

Discrete subaortic stenosis (DSS) is a rare, congenital heart disease characterized by formation of a fibrotic membrane in the subvalvular region of the left ventricular outflow tract (LVOT) of the heart, which then obstructs blood flow through the aortic valve .^1,2^ Up to 10% of patients with congenital heart disease present with a LVOT obstruction,^3^ and DSS accounts for up to 15% of these cases.^4^ When left untreated, the DSS membrane will continue growing and restrict blood flow through the LVOT, resulting in high rates of morbidity and mortality.^1,2,5^ Currently the only option for treatment is surgical removal of the membrane. Physicians typically wait for the membrane to progress into an “optimal surgical window,” which was defined in the 2008 ACC/AHA guidelines as peak instantaneous echocardiographic gradient greater than 50 mmHg, mean gradient greater than 30 mmHg, or catheter measurement of the resting peak-to-peak gradient greater than 50 mmHg.^1,3,6^ Following this surgical removal, however, up to 29% of DSS patients have recurrence of the membrane within 13 years of their first membrane resection.^1,5^

The underlying cellular mechanism that drives membrane formation in DSS is unknown; however, there are clues in the disease presentation. For example, the membrane is composed of fibrotic extracellular matrix proteins and DSS patients present with left ventricular hypertrophy, both hallmarks of cardiac fibrosis.^1,2^ Additionally, the aortoseptal angle is steeper than normal in patients with DSS, which could contribute to elevated wall shear stress within the LVOT.^1,2,7^ In the LVOT, physiological wall shear stresses range from 10-15 dynes/cm^2^, and DSS patients experience supra-physiological wall shear stresses that range from 30-55 dynes/cm^2^.^3,7,8^

The poor understanding of the mechanism of DSS is due in part to the lack of suitable in vitro models. To overcome this limitation, we created an in vitro model of DSS by investigating the effects of elevated wall shear stress levels on endocardial endothelial cells (EECs), the cells that line the LVOT. We isolated EECs from porcine hearts, exposed the cells to varying levels of shear stress using a cone-and-plate device,^9^ and then characterized the cellular inflammatory and fibrotic responses. This in vitro model provided a customized environment to understand how varied levels of shear stress impact the phenotype and cellular signaling mechanisms of the EECs, consistent with how they would be directly exposed to high shear stresses within the LVOT in DSS. This experimental design enabled us to distinguish how cellular responses are influenced by the magnitude of applied shear stress. We hypothesized that exposing EECs to pathological wall shear stress would initiate the endothelial-to-mesenchymal transition (EndMT) cell signaling pathway. The aim of this study was to determine the effects of time duration and magnitude of shear stress on EECs by characterizing cellular alignment, gene expression, and protein expression to elucidate the cellular signaling mechanisms that link elevated wall shear stress to the onset of DSS.

## Materials and Methods

### Endocardial endothelial cell isolation & culture

Endocardial endothelial cells were isolated from porcine hearts obtained from a local commercial abattoir (Animal Technologies, Tyler, TX) as previously described;^10^ the hearts of young adult pigs (6-12 months old) are commonly used as an experimental model for human hearts due to their anatomical and structural similarities and convenient availability.^11^ Cells were cultured on tissue culture treated flasks with Endothelial Growth Medium-2 (EGM-2, Lonza), which was changed every other day until the cells reached ∼75% confluence. For experiments, cells between passages 5-10 were plated in 60 mm tissue culture treated petri dishes (Nunc EasYDish, Thermo Fisher) at a density of 0.8×10^6^ cells per dish. The cells were grown to confluency prior to further experimentation.

### Applying shear stress with a cone-and-plate device

A cone-and-plate device was used to apply shear stress (**Supplemental Figure 1**), as previously described.^9^ The cone had a smooth surface with an angle of 0.5°. The shear stress applied by the cone was determined as 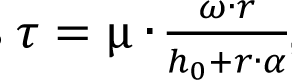, where μ = wall shear stress, µ = dynamic viscosity of fluid, ω = angular velocity of the cone, r = radius of the cone, h_0_= gap height, and α = angle between the cone-and-plate, as previously reported.^9^ After sterilizing the cones via UV light for at least 3 hours, the gap height of the cone was adjusted to 300 µm using the collar shaft. The adjusted cone was then loaded onto a petri dish containing EECs and 2.0 mL of EGM-2. The system was then placed on a magnetic stir plate with a controllable rotational speed (Cole-Parmer Modular Stirrer Control Unit and Magnetic Stirring Units) and the entire system was placed in a 5% CO_2_ incubator at 37°C. The shear stress levels used in experiments were static conditions (no shear stress), low shear (15 dynes/cm^2^), medium shear (20 dynes/cm^2^), or high shear (35 dynes/cm^2^). Clinical magnitudes of shear stress provided the basis for these experimental values. The low shear condition was based on physiological LVOT wall shear stresses, which range from 10-15 dynes/cm^2^, whereas the high shear condition was based on pathological LVOT wall shear stresses that range from 30-55 dynes/cm^2^ in DSS patients.^3,7,8^ To evaluate whether an intermediate, greater than physiological shear stress condition could trigger fibrotic signaling, we set a medium shear condition at 20 dynes/cm^2^. While the physiological environment of the LVOT involves complex flow patterns and DSS is characterized by flow disturbances, the aim of this study was to determine if the supra-physiological magnitude of shear stress would be sufficient to trigger the EndMT transition in EECs. Using the equation above, the desired shear stress was used to determine the required rotational setting of the stir plate (**Supplemental Table 1**). When starting the experiments, shear stress was added progressively in a ramped manner over a period of 5 minutes until the target magnitude of shear stress was achieved to avoid the abrupt application of high shear stress. The experimental overview provided in **Supplemental Table 2** shows the timepoints (1.5, 8, or 16 hrs) and shear conditions for this initial shear study.

### Cell imaging and analysis

Brightfield images of the cells were obtained before and after 16 hours of shearing, using a Nikon Ti2 Eclipse microscope. Representative images were reviewed for technical quality, and then selected to illustrate the most commonly observed morphology for each condition. After imaging, alignment analysis was performed on 10 representative non-overlapping images for each shear condition using ImageJ with the OrientationJ plugin. This plugin produced cell orientation data as well as images of EECs that were overlain with a color map corresponding to the degree of orientation.

### Quantitative polymerase chain reaction

Gene expression of key markers of cell phenotype, fibrosis, and inflammation were evaluated after 1.5, 8, or 16 hours of shearing. Total RNA was isolated from samples with TRIzol reagent (Thermo Fisher), then purified using the Direct-zol RNA MicroPrep Kit (Zymo Research) according to manufacturer’s instructions. RNA quality and quantity was evaluated using the NanoDrop 2000. RNA samples were stored at -80°C until further processing. cDNA was synthesized from RNA using the High Capacity cDNA Reverse Transcription Kit (Thermo Fisher), then stored at -20°C. Quantitative reverse transcriptase polymerase chain reaction (qRT-PCR) was performed using the iTaq Universal Probes Supermix (Bio-Rad) with Taqman porcine primers specific to *CD31*, *ACTA2*, *VCAM1*, *ICAM1*, and *GAPDH* (**Supplemental Table 3**). qPCR for *SNAI1* expression was performed using the RT^2^ SYBR Green qPCR Mastermix (Qiagen) with RT^2^ qPCR Primer Assay porcine primers specific to *SNAI1* and *GAPDH*. Fold changes were calculated using the ΔΔCT method with normalization to *GAPDH* and controls.

### Fibrosis profiler array

For a more comprehensive analysis of the fibrotic reponse of the EECs to shear stress, we utilized the RT^2^ Profiler PCR Array - Pig Fibrosis (Qiagen). RNA samples were isolated and purified as described above. cDNA was synthesized using the RT^2^ First Strand Kit (Qiagen). All kits were used according to manufacturer’s instructions and the array data was analyzed using the Qiagen GeneGlobe data analysis tool.

### Small molecule inhibition of Snail

To test the role of Snail in mechanotransduction signalling in EECs, a small molecule inhibitor of Snail, CYD19 (Aobious), was applied to cells. CYD19 inhibits Snail1 in vitro and in vivo with a post-translational modification that promotes degradation of the Snail1 protein through the ubiquitin-proteasome pathway.^12^ The molecule was dissolved in dimethylsulfoxide (DMSO) and stored at room temperature in a stock solution concentration of 4.0937 µM. CYD19 was added to the cells at a concentration of 50 nM, as recommended by a previous study.^12^ Cells were treated with CYD19 or the vehicle control (an equivalent volume of DMSO) for 2 days. The cell culture media was then changed and the cells were treated with static or high shear stress conditions for 1.5 or 24 hours. These timepoints were selected to capture acute gene expression (1.5 hours) and the subsequent protein expression (24 hours).

### Flow cytometry

Flow cytometry was performed to evaluate the EECs protein expression. EECs were rinsed with phosphate buffered saline (PBS) then lifted with Accutase (Sigma-Aldrich), transferred to a conical tube, and centrifuged for 5 minutes at 150 x*g*. The cells were then resuspended in 1% fetal bovine serum (FBS, Fisher Scientific) in PBS at a density of 1×10^6^ cells per 100 µL within multiple microcentrifuge tubes.

For extracellular protein analysis, the antibodies and corresponding isotype controls were added to their respective samples’ tubes at concentrations recommended by the manufacturer (**Supplemental Table 4**). The samples were protected from light, incubated at room temperature for 15 minutes, then washed twice with 1 mL of 1% FBS in PBS.

For intracellular protein analysis, the cells were fixed in 4% paraformaldehyde at 4°C for 20 minutes, then permeabilized in 1% Tween20, 1% FBS in PBS at room temperature for 15 minutes. The cells were then centrifuged and resuspended in 100 µL of the 1% Tween20, 1% FBS in PBS solution. The antibodies and corresponding isotype controls were then added (**Supplemental Table 4**). The samples were protected from light, incubated with the antibody at room temperature for 45 minutes, then washed with 1 mL of 1% FBS in PBS.

The stained cells were resuspended in 500 µL 1% FBS in PBS and transferred to polystyrene 5 mL flow cytometry tubes (MTC Bio). Flow cytometry was performed on a Sony MA900 flow cytometer, and analysis was performed in FlowJo Version 7.

### Statistical analysis

Statistical analysis was performed in GraphPad Prism Version 9.4.1. For most of the experiments, a one-way ANOVA test was performed, with post-hoc analysis performed using Tukey’s test. All data was graphed as mean±standard deviation. For the small molecule inhibitor experiments, a two-way ANOVA was performed, with post-hoc analysis performed using Tukey’s test. Results with p<0.05 were considered statistically significant. Comparisons with p-values between 0.05-0.15 were considered as non-significant trends.

## Results

### Shear stress drives EEC alignment

After 16 hours, brightfield images of the EECs were captured at multiple locations around the dish, and we analyzed the distribution of orientation of 10 samples for each shear condition. After shearing, the cells aligned in the direction of the fluid rotation (**Figure 1**). In the static condition, the cells had a non-uniform distribution of alignment. After applying low (15 dynes/cm^2^) or medium (20 dynes/cm^2^) magnitudes of shear stress, the cells aligned in a more uniform direction as shown by the dominant peak at approximately 45 degrees. Interestingly, in the high shear stress condition (35 dynes/cm^2^), the cells displayed a non-uniform alignment, similar to the static condition.

**Figure 1.**
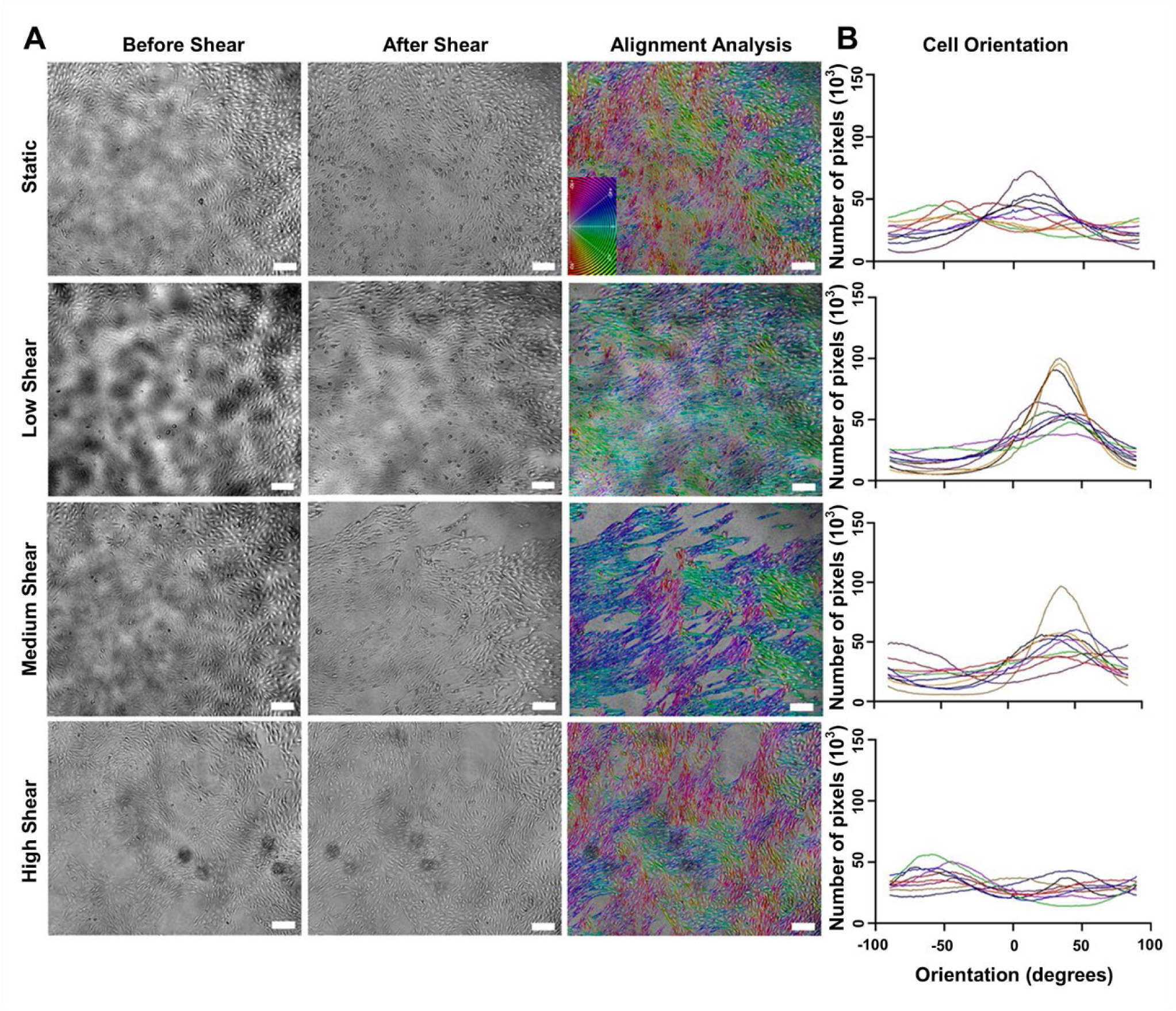
**(A)** Representative brightfield images of EECs before and after exposure to shear stress for 16 hours. The last column color maps the alignment of the EECs following exposure to shear. The color map code (inset of static condition) shows how the cell orientation corresponds to color. Scale bar = 200 µm. These were individual samples analyzed at the same location on the petri dish. **(B)** The cell orientation of 10 representative samples of imaged EECs for each shear condition. Each line represents an individual sample analyzed at the same location on the petri dish.

### Shear stress drives time dependent gene expression

We used qRT-PCR to analyze the gene expression of the EECs in response to varying shear rates over time (**Figure 2**), examining *CD31 and ACTA2* as endothelial and fibrotic markers, respectively, and *ICAM1* and *VCAM1* as inflammatory markers. These phenotypic markers were selected as representative based on previous studies evaluating endothelial fo mesenchymal transition in endothelial cells exposed to shear stress ^13–17^. The resulting cellular response was dependent on shear rate in a way that differed between time points. *CD31* gene expression remained largely unchanged across time points and shear conditions, however, *CD31* was downregulated in the low shear condition at 16 hours (37% decrease; p=0.0416) and a trend of downregulation of *CD31* was observed in the high shear condition at 1.5 hours (21% decrease; p=0.0818). Overall, the data indicates that the cells maintained an endothelial phenotype. *ACTA2*, which encodes for the protein α-smooth muscle actin, was downregulated in response to shear at all magnitudes, but only at the 8-hour time point (51-61% decrease; p<0.0001 for L, M, & H). The inflammatory marker, *ICAM1* was upregulated at 1.5 hours in response to increased shear (26-38% increase; p=0.0006-0.0096 for M & H), but did not demonstrate differences at later time points, other than a slight trend of a reduction with shear at 8 hours (15% decrease; p=0.0569). In contrast, *VCAM1* was downregulated by all shear magnitudes at 8 hours (44-57% decrease; p<0.0001 – p=0.0004). At 16 hours, a reduction in *VCAM1* expression was observed in the low and medium shear conditions (72-77% decrease; p=0.0003-0.0006), yet in the high shear condition the downregulation of VCAM1 was observed only as a slight trend (36% decrease; p=0.0681).

**Figure 2.**
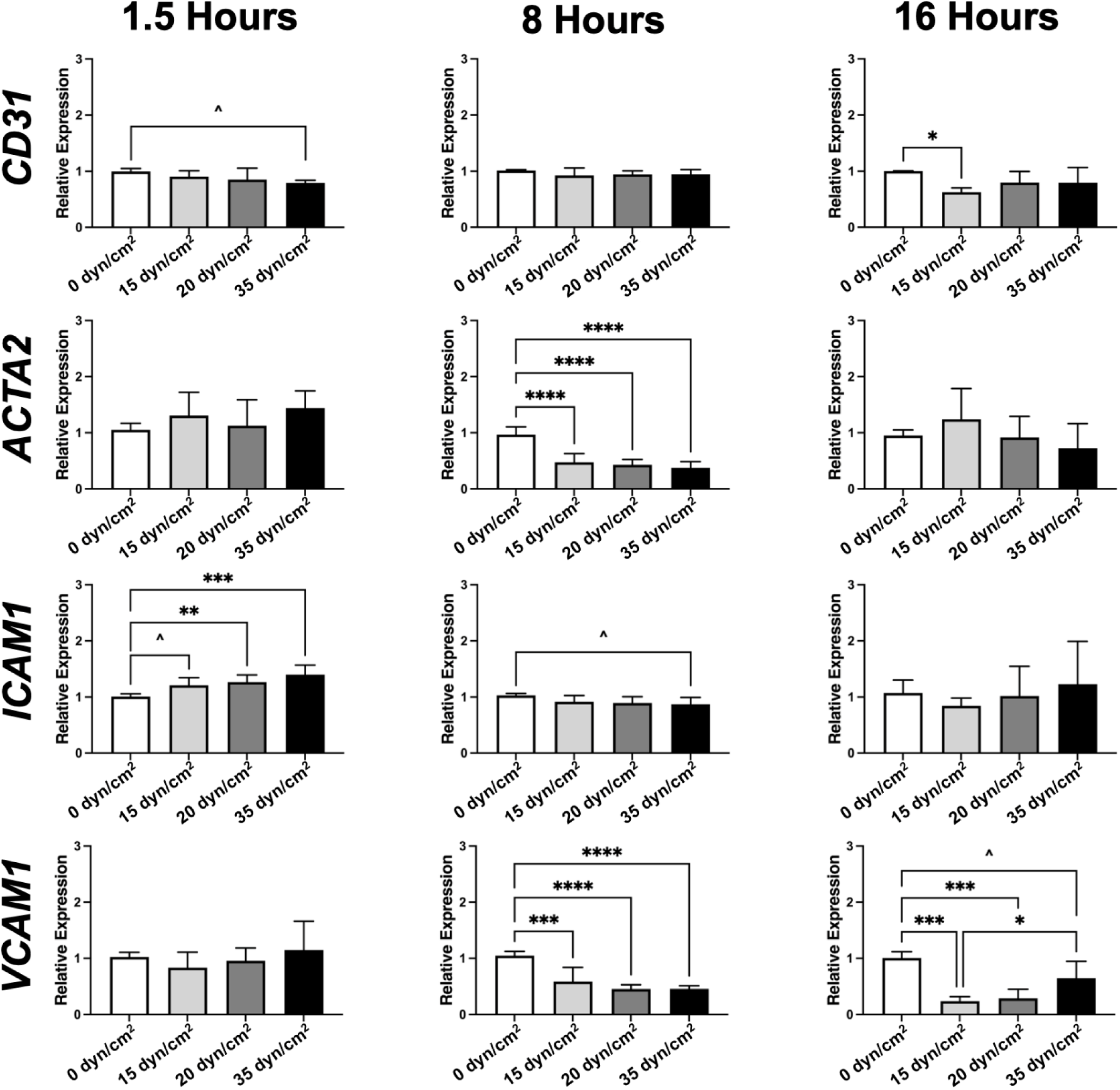
Gene expression of *CD31*, *ACTA2*, *ICAM1*, and *VCAM1* by EECS under static conditons and 15-35 dyn/cm^2^ conditions for 1.5, 8, and 16 hours. N=3. Data was normalized to the static condition for each time point, analyzed by one-way ANOVA, and expressed as mean ± SD. ^p=0.05-0.082, *p=0.01-0.05, **p=0.001-0.01, ***p=0.0001-0.001, ****p<0.0001.

### Fibrosis array suggests role for Snail in shear stimulated EECs

Based on these initial findings of cellular response to shear stress, we used a PCR profiler array to analyze expression of 84 fibrosis-related genes in either static or high shear stress (35 dynes/cm^2^) conditions at 3 time points (1.5 hours, 8 hours, and 16 hours). The focus on the high shear stress condition was based on pathological measurements in DSS ^7^ and on how this condition demonstrated the greatest changes in *ACTA2*, *ICAM1*, and *VCAM1* expression. Overall, the fibrosis array found that more genes were upregulated (68 genes across all time points) than downregulated (25 genes across all time points) in response to shear stress (**Supplemental Figure 2**). The heat map of the profiler array showed that there was consistency of gene expression within each experimental group, but that the expression profiles changed over time (**Figure 3, Supplemental Table 5**), supporting the PCR results.

**Figure 3.**
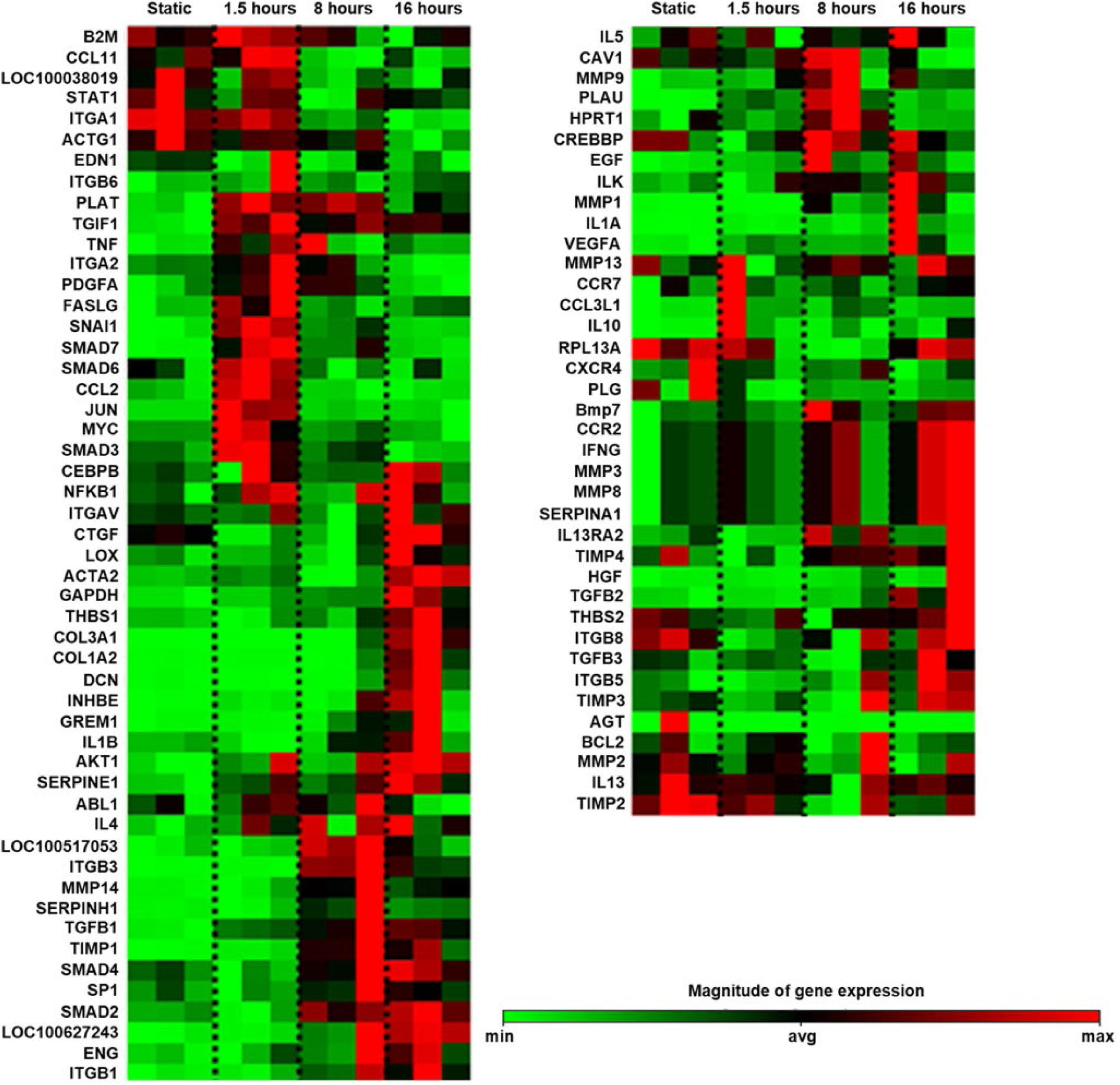
Fibrosis profiler array comparing the gene expression of static EECs to EECs exposed to high shear stress at 1.5, 8, and 16 hours, respectively. The samples are grouped by shear condition and time point. A separate z-distribution is shown for each gene. Green = minimum magnitude of gene expression; Red = maximum magnitude of gene expression. Refer to Supplemental Table 4 to view full data set.

The genes that were strongly upregulated or downregulated in response to shear stress provided insight into the involvement of specific signaling pathways. *SNAI1*, which encodes for the transcription factor Snail1, was very strongly upregulated (26.6 fold regulation (FR), p=0.0014) at the earliest time point of 1.5 hours, whereas ECM genes such as *COL3A1* (183.81 FR, p=0.0013), *COL1A2* (38.86 FR, p=0.0145), and *DCN* (90.73 FR, p=0.0315) were significantly upregulated at the later time point of 16 hours (**Table 1** and **Supplemental Table 5**). Several other genes relevant to ECM, cell-matrix adhesion, and fibrosis were also upregulated significantly for at least one of the later time points, including *ITGB3* (8.02-14.81 FR), *TFGB1* (2.76-2.93 FR), *TGFB2* (4.71 FR), *MMP14* (2.07-2.78 FR), and *TIMP1* (11.32-15.17 FR, all p<0.05). In contrast, selected signaling factors with an endothelial role, such as *CTGF* (−2.50 to -2.59 FR) and *CCL11* (−4.33 to −11.11 FR), were significantly downregulated (p<0.05) at different time points. Based on this data, we hypothesized that *SNAI1* was a mechanosensitive marker that was upregulated at early time points in response to shear. We speculated that this then initiated signaling pathways that lead to the increased ECM genes observed at later time points, which were indicative of EndMT.

**Table 1.**
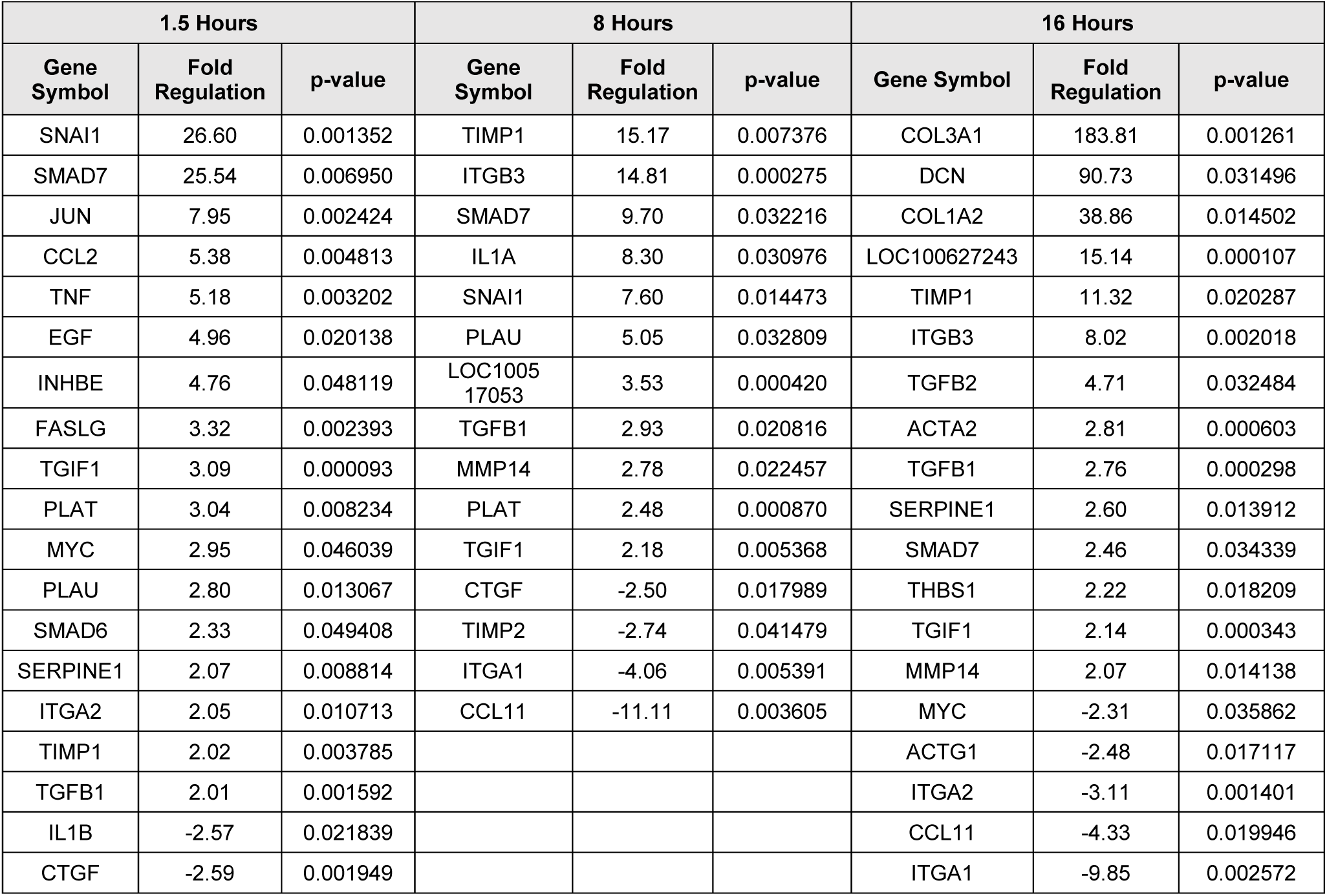
Significantly upregulated or downregulated fibrotic gene expression of EECs exposed to high shear stress (35 dynes/cm^2^) in comparison to static culture (0 dynes/cm^2^) for 1.5, 8, or 16 hours. Genes are listed in order of highest amount of fold upregulation to greatest amount of downregulation.

### Small molecule inhibition of Snail transiently influenced EndMT markers

To investigate this hypothesis, cells were treated with CYD19, which inhibits Snail1 via a post-translational modification,^12^ prior to the application of high shear stress. We observed an initial shift in cell morphology between the CYD19-treated and the vehicle-treated cells, as shown in **Figure 4A**. The vehicle-treated EECs had a very similar elongated and swirling morphology to the EECs shown in previous figures, whereas the CYD19-treated EECs became more round and cobblestone-like.

**Figure 4.**
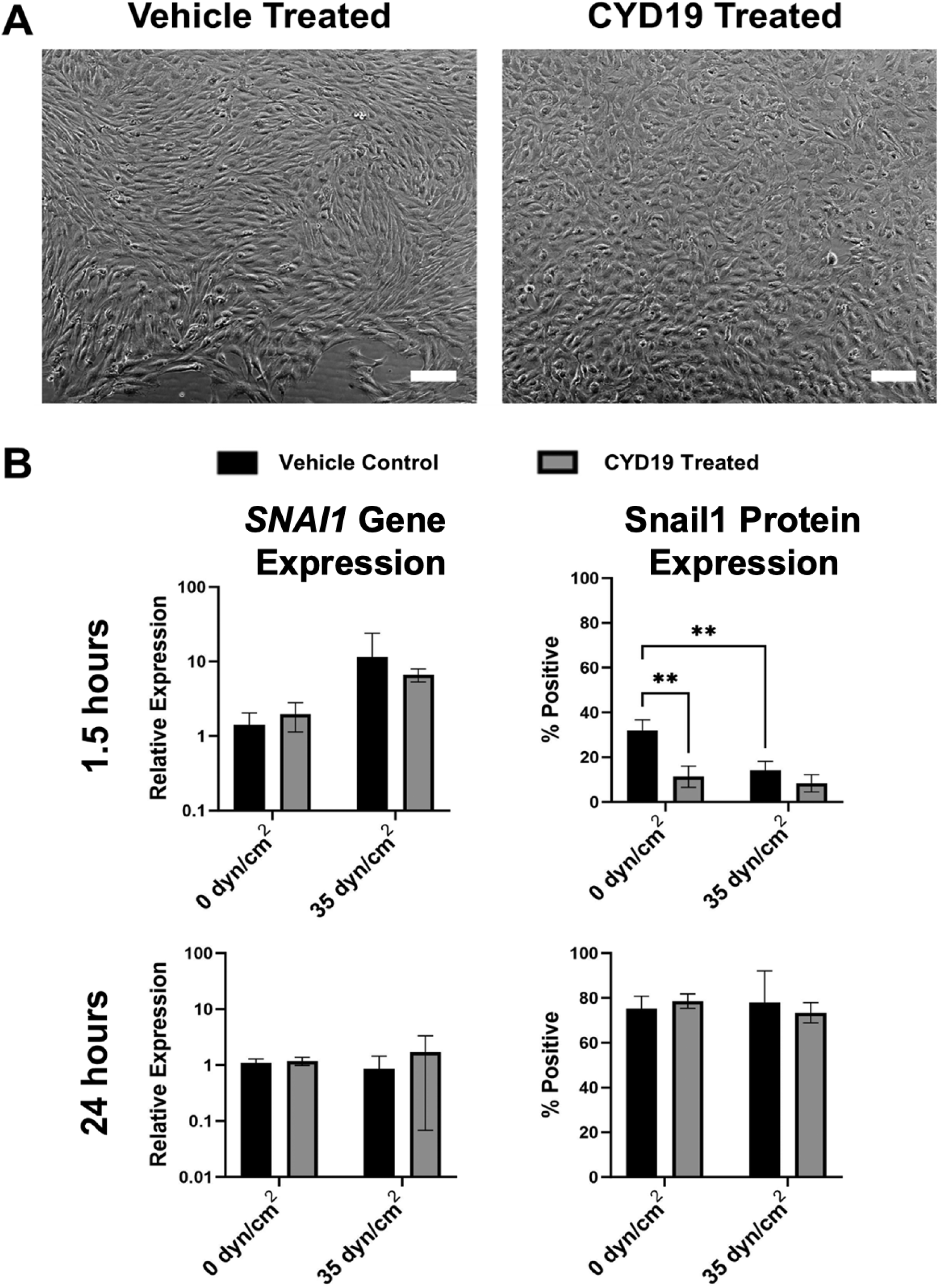
**(A)** Comparison of static vehicle-treated EECs (left) and static CYD19-treated EECs (right). Scale bar = 100 µm. **(B)** *SNAI1* gene and Snail1 protein expression from EECs cultured under either static conditions or high shear stress, and treated with either vehicle control or the small molecule inhibitor CYD19 at 1.5 hours and 24 hours. Data is expressed as mean ± SD. *p=0.01-0.05, **p=0.001-0.01, ***p=0.0001-0.001, ****p<0.0001.

At 1.5 hours after the application of shear stress, the gene expression of *SNAI1* appeared to be greater in the untreated group (**Figure 4B**), consistent with the fibrosis PCR assay data, although this difference was not statistically significant due to a high data spread. Treatment with CYD19 did not significantly change the gene expression of *SNAI1* in either of the static or high shear stress conditions, which was expected since CYD19 inhibits Snail1 on the protein level. The magnitude of difference between the static and high shear group appeared lessened in the CYD19 treated samples. In contrast with gene expression, at 1.5 hours Snail1 protein expression in the static cultured EECs was downregulated by 65% in response to CYD19 treatment compared to untreated (vehicle) controls (p=0.0019). Interestingly, at 1.5 hours protein levels of Snail1 were decreased by 56-74% in response to high shear stress in both groups (p=0.0008-0.0047). By 24 hours, *SNAI1* gene expression was not affected by either CYD19 treatment or the application of shear stress. Furthermore, Snail1 protein expression was abundant across all of the samples at the 24 hour time point (70-80%) compared to the expression level at 1.5 hours (10-40%).

At 1.5 hours, high shear stress reduced the protein expression of two mesenchymal phenotypic markers, α-smooth muscle actin (α-SMA) and vimentin (**Figure 5A**-**B**). At 1.5 hours, α-SMA protein was reduced in both the treated and untreated high shear groups by 62-63% (p=0.0399-0.0446) and vimentin protein was reduced reduced in both the treated and untreated high shear groups by 30-39% (p=0.0002-0.0012) when compared to their respective static untreated controls. At this timepoint, CYD19 treatment caused a slight reduction in vimentin protein for the statically cultured EECs (p=0.0540, trend). Otherwise, at 1.5 hours CYD19 treatment had no effect on protein expression for these two markers, and neither factor had a significant effect on *ACTA2* or *VIM* gene expression. At 24 hours, high shear stress downregulated *ACTA2* gene expression (71-86% decrease, p=0.0003-0.0012), α-SMA protein expression (42-61% decrease, p=0.0338-0.0043), and VIM gene expression (80-68% decrease, p<0.0001). Again, there was no significant effect of CYD19 treatment on expression levels, although there was a trend of reduced α-SMA protein in the statically cultured EECs (p=0.0709). It was noteworthy that, in all groups, more cells were positive for α-SMA protein at 24 hours (20-60%) than at 1.5 hours (1-3%). Similarly, vimentin protein expression was uniformly strong in all groups at 24 hours (<90%) compared to the protein expression levels at 1.5 hours (40-80%).

**Figure 5.**
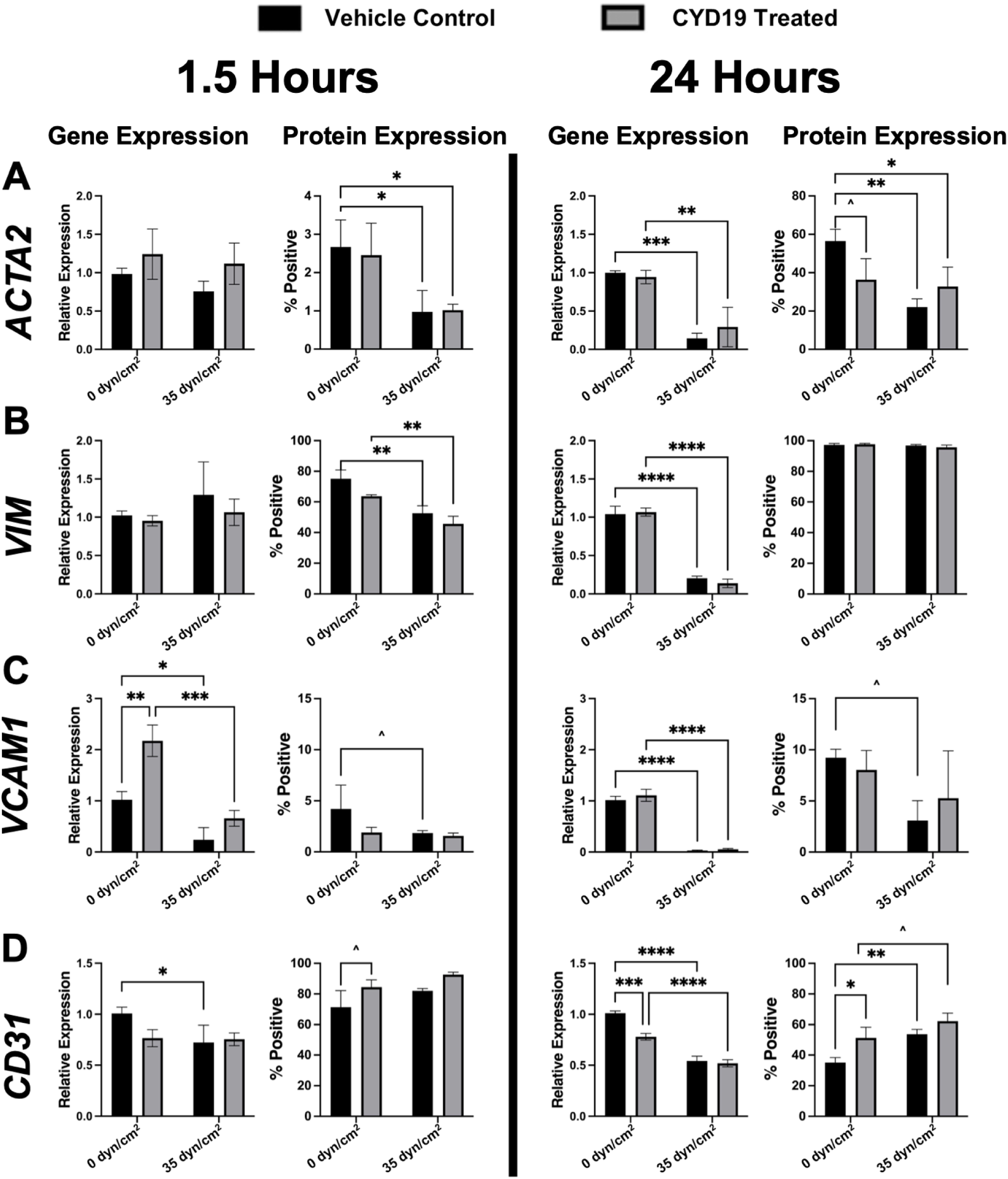
Gene and protein expression of **(A)** α-smooth muscle actin, **(B)** vimentin, **(C)** VCAM1, and **(D)** CD31 from EECs cultured under either static conditions (0 dyn/cm^2^) or high shear stress (35 dyn/cm^2^), and treated with either vehicle control or the small molecule inhibitor CYD19 at 1.5 hours and 24 hours. Data is expressed as mean ± SD. ^p<0.06, *p=0.01-0.05, **p=0.001-0.01, ***p=0.0001-0.001, ****p<0.0001.

In contrast to the mesenchymal markers, gene expression of the inflammatory marker *VCAM1* at 1.5 hours was upregulated 113% in response to CYD19 treatment at static conditions (p=0.0011), and was downregulated 70% in response to CYD19 treatment and high shear stress when compared to the CYD19 treated static condition (p=0.0002, **Figure 5C**). Additionally, gene expression of *VCAM1* at 1.5 hours was downregulated 76% in response to high shear stress (vehicle control) when compared to the static vehicle control (p=0.0114, **Figure 5C**). At 24 hours, *VCAM1* gene expression was downregulated 95-97% in response to high shear stress (p<0.0001), but not affected by CYD19. VCAM1 protein expression was not influenced by CYD19 treatment at either timepoint, but there were trends of reduced VCAM1 protein expression in the untreated groups following high shear stress for 1.5 hours (p=0.1563) and 24 hours (p=0.0908).

Finally, the gene expression of endothelial marker *CD31* at 1.5 hours was downregulated 28% in response to shear stress (p=0.0428, **Figure 5D**). At this timepoint, CYD19 did not affect *CD31* gene expression but did cause a slight non-significant trend of greater CD31 protein expression (p=0.1077). At 24 hours, *CD31* gene expression was downregulated 23% by CYD19 treatment (p=0.0002) and downregulated 49% by exposure to high shear stress (p<0.0001). The protein expression of CD31 at this later timepoint was upregulated by both the CYD19 treatment (46%, p=0.0175) and the application of high shear stress (Vehicle control: 53% increase, p=0.0079; CYD19 treatment: 22% increase, p=0.0975, trend). In contrast to the expression levels of mesenchymal protein markers between timepoints, CD31 protein expression was notably lower at the 24 hour time point (30-65%) than at the 1.5 hour time point (70-90%).

## Discussion

The altered subaortic anatomy found in DSS affects the local blood flow and consequently the magnitude of shear stresses^1^ that are experienced by the endocardial endothelial cells in the LVOT. In an effort to mimic the disease pathologies observed in DSS, we applied physiological and pathological levels of wall shear stress to EECs and characterized the cellular response through alignment, gene expression, and protein expression. The main findings of these experiments were that first, the EECs were mechanoresponsive and aligned in response to low and medium shear stress, but the collective response of EECs to high shear stress was more complex. Second, with increasing duration of exposure to high shear stresses, there was increased gene expression of many factors related to EndMT, especially those relevant to ECM remodeling, cell-matrix adhesion, and other mesenchymal markers. Third, the use of the CYD-19 inhibitor was effective in inhibition of Snail1 protein and mesenchymal transition, but the effects of this inihibitor were only transient and frequently overshadowed by the effects of high shear stress.

The EECs demonstrated mechanosensitivity by adjusting their alignment in response to shear, yet this realignment occurred in a shear magnitude-dependent manner. When comparing the low shear, medium shear, and high shear graphs, the cellular alignment is similar, with the most uniformity in cell alignment found in the medium shear condition. The shear-driven alignment of the EECs in low and medium shear rates is a phenomenon that has been observed in many studies where shear stress is applied to endothelial cells.^18,19^ However, after applying high shear stress to EECs, the condition most closely mimicking the disease state observed in DSS, the cells did not exhibit uniform alignment in response to the application of shear stress; we speculate that this demonstrates that the EECs were transitioning to a mesenchymal phenotype, such as a fibroblast or myofibroblast. Indeed, the decrease in uniformity of cellular alignment has been previously linked with cells beginning to lose their endothelial phenotype.^16,20^ The loss of alignment has been shown to increase inflammation,^16,20^ which is demonstrated by the significant increase of *ICAM1* gene expression at 1.5 hours in our study (**Figure 3**). Interestingly, *VCAM1*, an inflammatory marker, was downregulated with the application of shear stress at later timepoints. These *ICAM1* and *VCAM1* results, which are consistent with other investigations applying shear stresses to endothelial cells^16^ could be reflective of the early stage of EndMT.

The application of high shear stresses to the EECs resulted in the expression of many early- and late-stage genes involved in EndMT. During the EndMT process, the endothelial cells lose their endothelial characteristics, and begin to express mesenchymal markers.^13,14,21^ The transition is a continuum, and endothelial cells undergoing EndMT co-express both endothelial and mesenchymal markers,^20^ although this co-expression can vary depending on the pathological environment and cell type.^15,20,21^ In this study, a link between high shear stress and EndMT was shown throughout the culture duration. EECs exposed to 1.5 hours of high shear stress upregulated genes related to transcription factors, growth factors, cytokines, chemokines, and the formation of a transitional extracellular matrix (ECM).^15^ By the 16 hour timepoint, the EECs experiencing high shear stress showed strongly upregulated gene expression of mesenchymal markers, ECM, and matrix remodeling enzymes such as *ACTA2*, collagens 1 and 3 (*COL1A2* and *COL3A1*), decorin (*DCN*), *ITGB3*, *MMP14*, *TIMP1*, and *TGFβ1*, all markers of late EndMT processes.^15,20,21^ At the same timepoint, high shear stress caused down-regulation of vascular endothelial markers such as *EDN1*. This effect of time, as well as shear stress magnitude, on the response of the EECs was evident in the initial PCR analysis as well as the fibrosis gene array. The high shear stress also caused a dramatic increase in expression of the transcription factor Snail1, an important regulator of EndMT.^13,16,22–25^ Therefore, we sought to determine if blocking Snail1 could inhibit development of EndMT in EECs using CYD19, a small molecule inhibitor of Snail1 protein post-translational modification.^12^

Inhibition of Snail1 post-translational modification was able to dampen EndMT by EECs transiently. Since the inhibitor affected the Snail1 protein, the strongest effect on Snail1 expression was observed at the protein level, although this was only at the early time point. At this time (1.5 hours), the Snail1 inhibition maintained a more endothelial, less mesenchymal phenotype in the statically cultured EECs as shown by trends of reduced vimentin and increased CD31 and a strong, significant increase in *VCAM1*. By 24 hours, the inhibitor had no effect on Snail1 expression at either the gene or protein level. Other than a trend of slightly reduced α-SMA protein expression in the CYD19 treated group, which may be a downstream remnant of the Snail1 inhibition, there was no effect of CYD19 treatment on the mesenchymal or inflammatory genes at 24 hours. In contrast, *CD31* expression was affected by CYD19 treatment at 24 hours, with slightly lower gene expression but slightly greater protein expression. When the magnitudes of protein expression were considered collectively, the EECs at 24 hours appeared more mesenchymal and less endothelial in their phenotypic characteristics, especially in the statically cultured groups. Thus, the CYD19 small molecular inhibition of Snail1 activity and EndMT appeared to be successful, but this effect was only broadly evident for a limited time.

Interestingly, the effects of high shear stress were much more pronounced than that of the small molecule Snail1 inhibitor. This effect of high shear was particularly evident for *VCAM1* expression at both time points, as well as early protein expression and late gene expression for both α-SMA and vimentin, regardless of Snail1 inhibition. These *VCAM1* and *ACTA2* responses were consistent with the 8 hour EEC responses to shear stress in the initial PCR study. These mechanobiological trends highlight the importance of investigating shear stress when modeling diseases of the heart and cardiovascular system, and also emphasize the challenges of inhibiting the complex cell signaling pathways that drive DSS formation.

This is the first study to apply supra-physiological levels of wall shear stress to EECs and characterize their response as related to DSS formation. EECs have been implicated in several cardiac diseases, yet shear stress studies have focused primarily on modeling the behavior of HUVECs in response to pathological shear rates.^25,26^ EECs have a unique endothelial phenotype, as shown through proteomic analyses documenting that EECs express a different phenotype from aortic endothelial cells and even vary their phenotype throughout the regions of the heart.^26,27^ It is well understood that abnormal shear stress induces EndMT in HUVECs and in aortic valve endothelial cells, and may contribute to vascular disease formation.^15,16,20,28–31^ However, additional studies are needed to gain further insight into the EEC response to pathological shear stress. In a recent study investigating the role of shear stress in the formation of endocardial fibroelastosis (EFE),^25^ EECs exposed to abnormal shear conditions (uniform laminar shear stress 5 dynes/cm^2^) showed signs of undergoing EndMT through the co-expression of endothelial and mesenchymal markers and the increased expression of Snail1, which they concluded contributes to EFE. This important work emphasizes that the specific phenotype and resulting signaling pathways of EECs are unique.

These results have provided novel insights into the EEC-specific responses to physiological and pathological levels of shear stress, but they should be considered in the appropriate experimental context. Although we used the recommended dose of CYD19, it will be important in the future to investigate dose dependent effects of the inhibitors of EndMT signaling.^12^ It is also likely that to recapitulate the full EndMT signaling process that accompanies DSS, additional cell types (such as cardiac fibroblasts and immune cells) would need to be included in the disease model. Additionally, while we investigated the cellular response to shear stress between 1.5 hours and 24 hours, it would be worthwhile to extend these experiments to longer time points to understand the time dependent nature of EndMT progression. This experimental time frame was intended to characterize the acute initial and early progression of EndMT in EECs exposed to supra-physiological magnitudes of shear stress. Conducting a study of 48 hours or up to 5 days would provide insights regarding whether the EndMT phenotypic transition would continue to progress to mesenchymal cells and would assess the stability of the phenotypic markers at extended timepoints. Clearly, further investigation is required to uncover the complex signaling mechanisms of the fibrotic membrane formation observed in DSS.

## Conclusions

In summary, this study demonstrated that endocardial endothelial cells are highly sensitive to shear stresses and respond by changing their alignment and their gene expression of key markers of cell phenotype, EndMT, and inflammation. Our findings are consistent with the hypothesis that pathological wall shear stress initates the EndMT cell signaling pathway. Although inhibition of Snail1 with CYD19 had a modest inhibitory effect on EndMT and caused a significant increase in *VCAM1* expression, this effect was largely limited to the earlier timepoint of 1.5 hours. These results establish a cellular mechanism in which acute exposure to pathological wall shear stress induces EndMT in EECs. This study provides an important characterization of the EEC response to elevated wall shear stress, and lays the groundwork for additional studies to elucidate how shear stress and EndMT contribute to the role of EECs in DSS and other LVOT pathologies.

## Declarations

### Ethics approval and consent to participate

Not applicable.

### Consent for publication

All authors have provided consent to publish this article.

### Availability of data and material

The corresponding author will make data available upon request.

### Authors’ contributions

Kathleen Brown, Sundeep Keswani, and Jane Grande-Allen led the study conception and design, with important contributions from Ravi Birla, Philippe Sucosky, and Jennifer Connell. Research funding was obtained by Jane Grande-Allen, Sundeep Keswani, and Kathleen Brown. Experimental preparation, data collection, and data analysis were performed by Kathleen Brown, Kim Phan, Tasneem Mustafa, Elysa Jui, and Fariha Ahmad. The first draft of the manuscript was written by Kathleen Brown. Ravi Birla, Jennifer Connell, Sundeep Keswani, and Jane Grande-Allen commented on previous versions of the manuscript. All authors read and approved the final manuscript.

## Acknowledgements

The authors thank Dr. Monica Fahrenholtz for her critical review of the manuscript.

