## Supplemental Materials for "Shear Stress Initiates Endothelial to Mesenchymal Transition in Endocardial Endothelial Cells"

### SUPPLEMENTARY MATERIALS

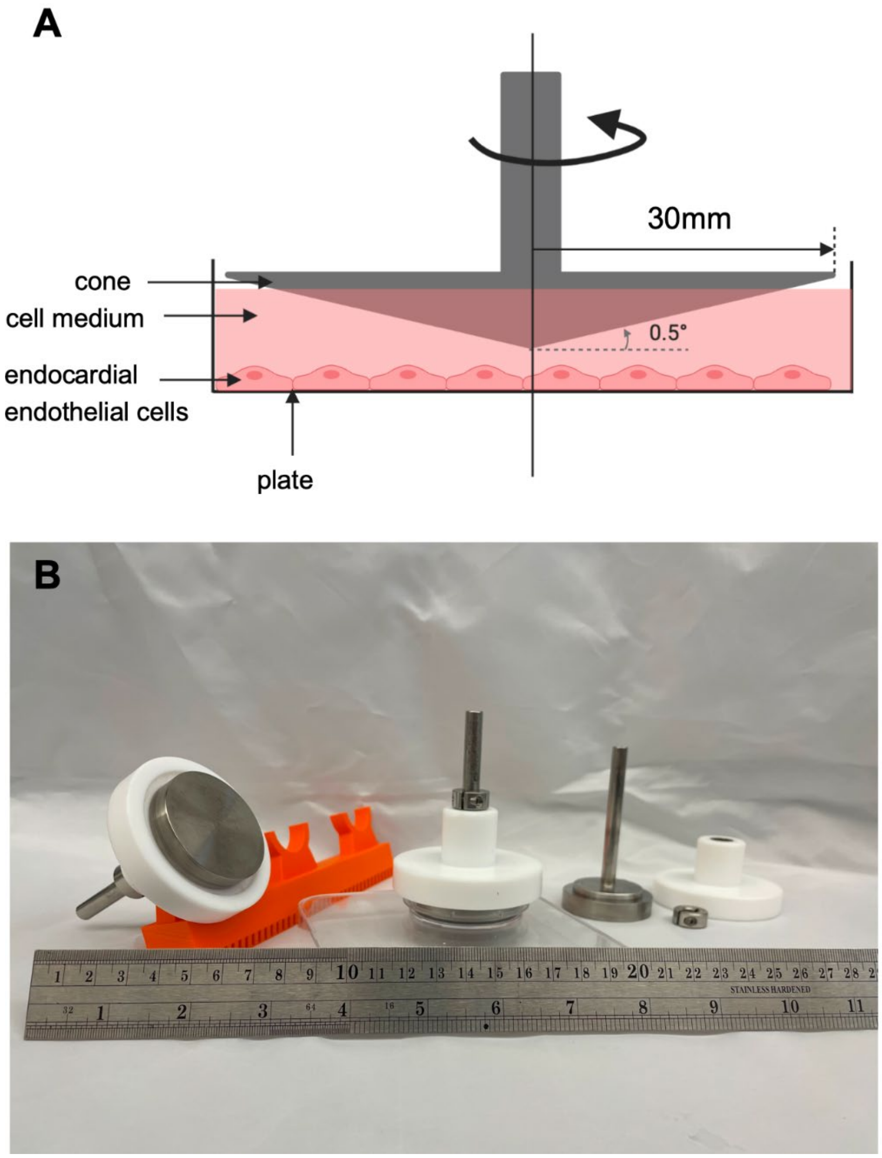


**Supplemental Figure 1.** Cone-and-plate Device. **(A)** Illustration of the cone-and-plate device. Endocardial endothelial cells are grown in a monolayer in a 60 mm petri dish, and the cone is loaded onto the petri dish containing the cells and cell culture medium and rotated using a controllable magnetic stir plate. **(B)** Photo of the cone and Teflon cover setup used in the shear stress studies. On the left of the photo, the cone is supported by a 3D printed holding device, which is used to position multiple cones towards a UV light for sterilization prior to use. In the center of the photo, the cone is loaded onto a petri dish. On the right of the photo, the cone, Teflon cover, and shaft collar are disassembled to show the individual parts used in the setup.

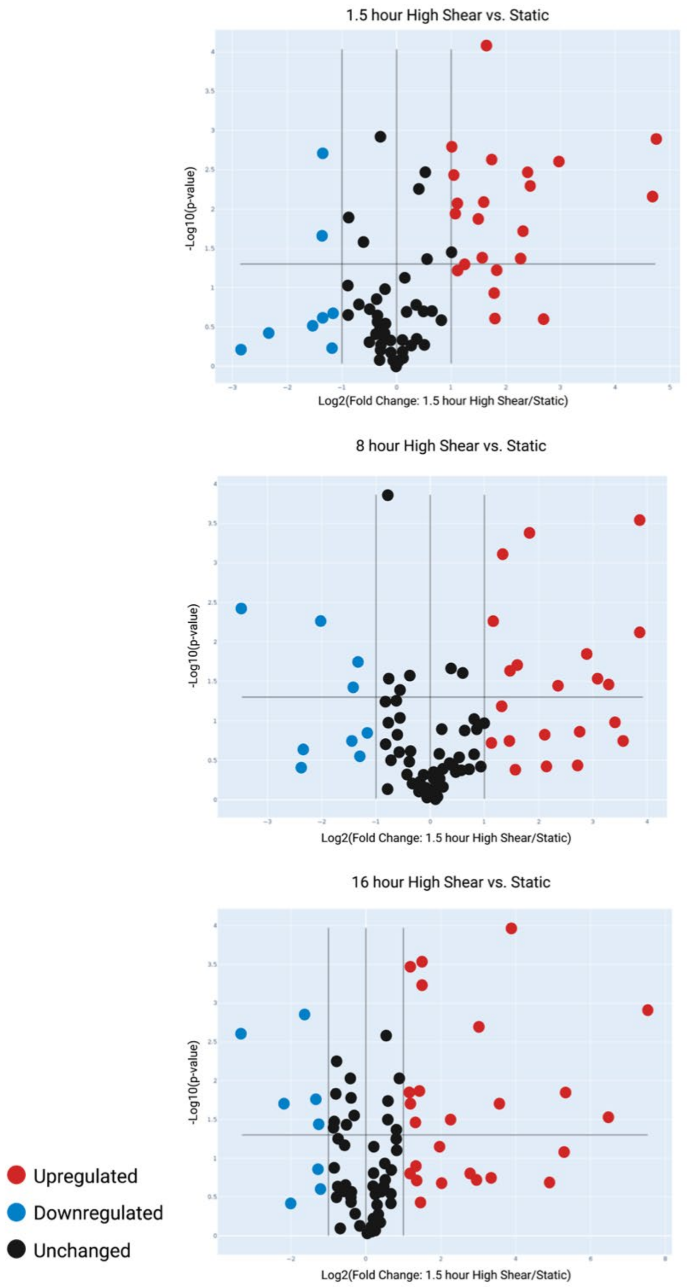


**Supplemental Figure 2.** Volcano plots comparing the gene expression of static EECs to EECs exposed to high shear stress at 1.5, 8, and 16 hours, respectively. Genes that are upregulated are shown as red dots, genes that are downregulated are shown as blue dots, and genes that remain unchanged are shown as black dots.

**Supplemental Table 1.** Cone-and-plate device settings.

| **Rotational Speed Setting** (RPM) | **Shear Stress Applied** (dynes/cm$^2$) |
|---|---|
| 280 | 15 |
| 370 | 20 |
| 660 | 35 |

**Supplemental Table 2.** Experimental conditions for shear stress experiments.

| | **Static Control** | **Low Shear Stress** (15 dynes/cm$^2$) | **Medium Shear Stress** (20 dynes/cm$^2$) | **High Shear Stress** (35 dynes/cm$^2$) |
|---|---|---|---|---|
| **1.5 hours** | Static 1.5 hours | Low shear 1.5 hours | Medium shear 1.5 hours | High shear 1.5 hours |
| **8 hours** | Static 8 hours | Low shear 8 hours | Medium shear 8 hours | High shear 8 hours |
| **16 hours** | Static 16 hours | Low shear 16 hours | Medium shear 16 hours | High shear 16 hours |

**Supplemental Table 3.** Taqman or Qiagen porcine primers specific to the genes of interest for PCR.

| **Gene of Interest** | **Primer Probe ID (Taqman) or GeneGlobe ID (Qiagen)** |
|---|---|
| *CD31* | Ss03392600_u1 (Taqman) |
| *ACTA2* | Ss04245588_m1 (Taqman) |
| *VCAM1* | Ss03390912_m1 (Taqman) |
| *ICAM1* | Ss03392385_m1 (Taqman) |
| *GAPDH* | Ss03374854_g1 (Taqman) |
| *SNAI1* | PPS07880A (Qiagen) |
| *GAPDH* | PPS00192A (Qiagen) |

**Supplemental Table 4.** Antibody information and concentrations used for flow cytometry.

| Antibody Information | Amount of Antibody Added to 100 μL Sample |
|---|---|
| **Extracellular Proteins** | |
| CD31 Monoclonal Antibody (LCI-4) [APC] – Thermo Fisher, #MA5-16769 | 10 μL |
| Mouse IgG1 kappa Isotype Control [APC] – Thermo Fisher, #17-4714-42 | 5 μL |
| VCAM-1 Monoclonal Antibody (1.G11B1) [FITC] – Thermo Fisher, #MA5-16430 | 4 μL |
| Igg1, K Mouse, Clone: P3.6.2.8.1, Isotype Control [FITC] – Thermo Fisher, #11-4714-42 | 4 μL |
| CD62E/CD62P Monoclonal Antibody (1.2B6) [PE] – Thermo Fisher, #MA5-28649 | 4 μL |
| Mouse IgG1 Isotype Control [PE] – Thermo Fisher, #MA518097 | 4 μL |
| **Intracellular Proteins** | |
| Snail Antibody [AlexaFluor® 750] – Novus Biologicals, # NBP2-27293AF750 | 4 μL |
| Rabbit IgG Isotype Control [AlexaFluor 750] – Novus Biologicals, # NBP2-24891AF750 | 4 μL |
| Vimentin Antibody (RV203) [PerCP] – Novus Biologicals, # NBP1-97671PCP | 4 μL |
| Mouse IgG1 Isotype control [PerCP] – Novus Biologicals, # IC002C | 4 μL |
| α-Smooth Muscle Actin Antibody [AlexaFluor 405] – Novus Biologicals, # NBP2-34522AF405 | 4 μL |
| Mouse IgG2a Isotype Control (M2AK) [AlexaFluor 405] – Novus Biologicals, # NBP1-96981AF405 | 4 μL |

**Supplemental Table 5.** Full data set of EEC gene expression and fold regulation in response to high shear stress (35 dynes/cm$^2$) at 1.5 hours, 8 hours, and 16 hours, respresented as a heatmap in Figure 4. In the Gene Symbol columns, the * symbol indicates that the gene's average threshold cycle is relatively high (>30) in either the control or the test sample, and is reasonably low in the other sample (<30), suggesting that the actual fold regulation value is at least as large as the reported fold regulation value. In the Fold Regulation columns, bolded numbers indicate that this gene exceeded the fold regulation threshold of 2, meaning this gene was upregulated (if positive value) or downregulated (negative value) in response to shear stress at this time point. In the p-value columns, bolded p-values indicate the gene expression observed is statistically significant (p<0.05) in comparison to the static control group. Genes are listed in order of the profiler array plate position. Note that genes for which the average threshold cycle was determined to be high (> 30) have been excluded from this table to ensure accuracy of reported results.

| **1.5 Hours** | | | **8 Hours** | | | **16 Hours** | | |
|---|---|---|---|---|---|---|---|---|
| **Gene Symbol** | **Fold Regulation** | **p-value** | **Gene Symbol** | **Fold Regulation** | **p-value** | **Gene Symbol** | **Fold Regulation** | **p-value** |
| ABL1 | 1.21 | 0.540121 | ABL1 | 1.15 | 0.695620 | ABL1* | -1.47 | 0.231918 |
| ACTA2 | 1.31 | **0.005508** | ACTA2 | -1.53 | 0.152235 | ACTA2 | **2.81** | **0.000603** |
| AKT1 | 1.11 | 0.075017 | AKT1 | -1.08 | 0.499801 | AKT1 | 1.14 | 0.071425 |
| BCL2 | -1.02 | 0.802490 | BCL2 | -1.27 | 0.588542 | BCL2* | -1.37 | 0.271104 |
| CAV1 | -1.17 | 0.104922 | CAV1 | -1.12 | 0.777320 | CAV1 | -1.87 | **0.014893** |
| CCL2 | **5.38** | **0.004813** | CCL11 | **-11.11** | **0.003605** | CCL11 | **-4.33** | **0.019946** |
| CTGF | **-2.59** | **0.001949** | CCL2 | -1.78 | 0.192672 | CCL2 | -1.92 | 0.135904 |
| CEBPB* | **-7.25** | 0.607630 | CTGF | **-2.50** | **0.017989** | CTGF | 1.16 | 0.270771 |
| JUN | **7.95** | **0.002424** | CEBPB | -1.30 | 0.242663 | CEBPB | 1.51 | 0.290460 |
| COL1A2 | 1.08 | 0.649424 | JUN | 1.00 | 0.935931 | JUN | -1.50 | 0.254432 |
| COL3A1 | 1.47 | **0.046892** | COL1A2 | 1.36 | 0.429572 | COL1A2 | **38.86** | **0.014502** |
| CXCR4 | -1.40 | 0.505999 | CXCR4 | -1.33 | 0.485514 | COL3A1* | **183.81** | **0.001261** |
| EDN1 | -1.23 | 0.604289 | EDN1 | -1.58 | 0.055340 | CXCR4 | **-2.25** | 0.250346 |
| EGF | **4.96** | **0.020138** | ENG | 1.45 | 0.319121 | DCN | **90.73** | **0.031496** |
| ENG | 1.10 | 0.609845 | IL1A | **8.30** | **0.030976** | EDN1 | -1.73 | **0.005902** |
| FASLG | **3.32** | **0.002393** | ILK | 1.03 | 0.557597 | ENG | 1.83 | **0.046697** |
| IL1B | **-2.57** | **0.021839** | ITGA1* | **-4.06** | **0.005391** | ILK | 1.13 | 0.226817 |
| ILK | -1.04 | 0.674167 | ITGA2 | 1.12 | 0.542976 | ITGA1* | **-9.85** | **0.002572** |
| INHBE | **4.76** | **0.048119** | LOC1005 17053 | **3.53** | **0.000420** | ITGA2* | **-3.11** | **0.001401** |
| ITGA1 | -1.14 | 0.470250 | ITGAV | -1.50 | **0.044684** | LOC10051 7053 | 1.55 | 0.154126 |
| ITGA2 | **2.05** | **0.010713** | ITGB1 | 1.56 | 0.130197 | ITGAV | 1.03 | 0.787618 |
| LOC10051 7053 | -1.01 | 0.926509 | ITGB3* | **14.81** | **0.000275** | ITGB1 | 1.80 | **0.047030** |
| ITGAV | -1.01 | 0.872866 | ITGB5 | -1.01 | 0.962301 | ITGB3* | **8.02** | **0.002018** |
| ITGB1 | 1.02 | 0.791480 | ITGB8 | -1.68 | 0.105258 | ITGB5 | 1.37 | 0.239532 |

| 1.5 Hours | | | 8 Hours | | | 16 Hours | | |
|---|---|---|---|---|---|---|---|---|
| **Gene Symbol** | **Fold Regulation** | **p-value** | **Gene Symbol** | **Fold Regulation** | **p-value** | **Gene Symbol** | **Fold Regulation** | **p-value** |
| ITGB5 | -1.19 | 0.301955 | LOC1000 38019 | -1.72 | **0.028793** | ITGB8 | -1.32 | 0.328220 |
| ITGB8 | -1.91 | **0.012929** | SNAI1* | **7.60** | **0.014473** | LOC10003 8019 | -1.69 | 0.050603 |
| LOC10003 8019 | -1.16 | 0.424999 | LOX | -1.31 | 0.327202 | LOX | 1.45 | 0.126836 |
| SNAI1* | **26.60** | **0.001352** | LOC1006 27243* | **6.86** | 0.145521 | LOC10062 7243* | **15.14** | **0.000107** |
| LOX | -1.04 | 0.678399 | MMP14 | **2.78** | **0.022457** | MMP14 | **2.07** | **0.014138** |
| MMP14 | 1.09 | 0.488623 | MMP2 | -1.49 | 0.259701 | MMP2 | -1.65 | 0.240076 |
| MMP2 | -1.22 | 0.268772 | MYC | -1.07 | 0.692770 | MYC | **-2.31** | **0.035862** |
| MYC | **2.95** | **0.046039** | NFKB1 | -1.10 | 0.671565 | NFKB1 | 1.01 | 0.991373 |
| NFKB1 | 1.27 | 0.173665 | PDGFA | 1.11 | 0.283864 | PDGFA | -1.32 | **0.009820** |
| PDGFA | 1.44 | **0.003379** | PLAT | **2.48** | **0.000870** | PLAT | 1.41 | 0.198602 |
| PLAT | **3.04** | **0.008234** | PLAU* | **5.05** | **0.032809** | PLAU | 1.52 | **0.030341** |
| PLAU | **2.80** | **0.013067** | SERPINE 1 | 1.75 | 0.097237 | SERPINE1 | **2.60** | **0.013912** |
| SERPINE1 | **2.07** | **0.008814** | SERPINH 1 | **2.48** | 0.061137 | SERPINH 1 | 1.52 | **0.019464** |
| SERPINH1 | 1.01 | 0.831593 | SMAD2 | 1.31 | **0.021648** | SMAD2 | 1.44 | **0.002571** |
| SMAD2 | -1.24 | **0.001113** | SMAD3 | -1.02 | 0.848107 | SMAD3 | -1.97 | **0.040385** |
| SMAD3 | **2.11** | 0.055446 | SMAD4 | 1.07 | 0.611947 | SMAD4 | 1.17 | 0.155932 |
| SMAD4 | -1.29 | 0.140945 | SMAD6 | -1.62 | 0.294007 | SMAD6 | -1.71 | 0.301776 |
| SMAD6 | **2.33** | **0.049408** | SMAD7 | **9.70** | **0.032216** | SMAD7 | **2.46** | **0.034339** |
| SMAD7 | **25.54** | **0.006950** | SP1 | 1.21 | 0.413718 | SP1 | 1.09 | 0.639097 |
| SP1 | -1.26 | 0.292278 | STAT1 | -1.72 | 0.056513 | STAT1 | -1.48 | 0.063858 |
| STAT1 | -1.19 | 0.360376 | TGFB1 | **2.93** | **0.020816** | TGFB1 | **2.76** | **0.000298** |
| TGFB1 | **2.01** | **0.001592** | TGFB2 | 1.42 | 0.405871 | TGFB2 | **4.71** | **0.032484** |
| TGFB2 | -1.03 | 0.791053 | TGIF1 | **2.18** | **0.005368** | TGIF1 | **2.14** | **0.000343** |
| TGIF1 | **3.09** | **0.000093** | THBS1 | 1.07 | 0.661091 | THBS1 | **2.22** | **0.018209** |
| THBS1 | -1.27 | 0.231704 | TIMP1 | **15.17** | **0.007376** | TIMP1 | **11.32** | **0.020287** |
| TIMP1 | **2.02** | **0.003785** | TIMP2 | **-2.74** | **0.041479** | TIMP2 | -1.94 | **0.034846** |
| TIMP2 | -1.38 | 0.190410 | TIMP3 | -1.75 | 0.713563 | TIMP3 | 1.24 | 0.383076 |
| TIMP3 | -1.53 | **0.028974** | VEGFA | 1.49 | **0.024391** | VEGFA | **2.51** | 0.187627 |
| TNF | **5.18** | **0.003202** | ACTG1 | -1.48 | 0.090930 | ACTG1 | **-2.48** | **0.017117** |
| VEGFA | 1.99 | **0.034918** | B2M | -1.32 | **0.028227** | B2M | -1.40 | **0.036057** |
| ACTG1 | -1.22 | 0.283476 | GAPDH | 1.15 | 0.129160 | GAPDH | 1.97 | **0.009484** |
| B2M | 1.02 | 0.777534 | HPRT1 | 1.05 | 0.475802 | HPRT1 | -1.31 | **0.016557** |
| GAPDH | 1.12 | 0.203054 | RPL13A | -1.75 | **0.000138** | RPL13A | -1.29 | **0.030081** |
| HPRT1 | -1.07 | 0.467381 | | | | | | |
| RPL13A | -1.22 | 0.346220 | | | | | | |
